# Methyl-SNP-seq reveals dual readouts of methylome and variome at molecule resolution

**DOI:** 10.1101/2022.06.28.498010

**Authors:** Bo Yan, Duan Wang, Romualdas Vaisvila, Zhiyi Sun, Laurence Ettwiller

## Abstract

Covalent modifications of genomic DNA are crucial for most organisms to survive. Amplicon-based high throughput sequencing technologies erase all DNA modifications to retain only sequence information for the four canonical nucleobases, necessitating specialized technologies for ascertaining epigenetic information. To also capture base modification information, we developed Methyl-SNP-seq, a technology that takes advantage of the complementarity of the double helix to extract the methylation and original sequence information from a single DNA molecule. More specifically, Methyl-SNP-seq uses bisulfite conversion of one of the strands to identify cytosine methylation while retaining the sequence of the other strand. As both strands are locked together to link the dual readouts on a single paired-end read, Methyl-SNP-seq allows detecting methylation status of any DNA even without a reference genome. Because one of the strands retains the original 4 nucleotide composition, Methyl-SNP-seq can also be used in conjunction with standard sequence-specific probes for targeted enrichment and amplification. We demonstrate the usefulness of this technology in a broad spectrum of applications ranging from allele-specific methylation analysis in humans to identification of methyltransferase specificity in complex bacterial communities.

## Introduction

The covalent modification of cytosine by a methyl group leads to the formation of 5-methylcytosine (5mC), a key epigenetic modification of genomic DNA that occurs in a large number of organisms and represents so far the best characterized form of DNA modification. In mammals, patterns of methylation are established early during embryogenesis and include X-chromosome inactivation, imprinting, and the repression of repeats and transposable elements (Greenberg and Bourc’his 2019). Not surprisingly, global or regional changes of DNA methylation are among the earliest events known to occur in cancer (Baylin and Jones 2016). Identifying methylation profiles in humans is a key step in studying disease processes and is increasingly used for diagnostic purposes.

In prokaryotes, the vast majority of genomes contain methylated nucleotides (Blow et al. 2016). Contrary to eukaryotes where the methylation sites are variable and subject to epigenetic states, bacterial methylations tend to be constitutively present at specific sites across the genome. These sites are defined by the methyltransferase specificity and, in the case of RM systems, tend to be fully methylated to avoid cuts by the cognate restriction enzyme. Current high throughput techniques for the identification of 5mC are performed by converting cytosine (C) to uracil (which is read as T during sequencing) leaving 5’methylcytosine (5mC) intact. This conversion is done using chemical treatment (bisulfite) or enzymatic treatment (EM-seq). In both methods, this conversion must be complete, leading to the loss of complementarity and separation of the two strands with thymine (T) being either a genuine T or the product of amplification after deamination of C. Consequently, separate experiments are needed to obtain accurate sequence variants and DNA modification information. Recently, a new technique has been developed that locks Watson and Crick strands together by a hairpin adapter followed by bisulfite treatment (Liang et al. 2021). However, because both strands are subjected to conversion, neither of the strands retains the original sequence information.

To circumvent these limitations, we developed Methyl-SNP-seq. Similar to the hairpin adaptor method previously published (Laird et al. 2004 Liang et al. 2021), we lock the forward and reverse strand together. The innovation is on our new design that permits a dual direct readout of the sequence and the methylation on the same paired-end read (see **Figure 1** for principle). So far, this dual readout was only possible with single molecule sequencing platforms such as Nanopore (Rand et al. 2017; Simpson et al. 2017) and PacBio (Clark et al. 2013). While single molecule sequencing requires the original DNA molecule to be sequenced in order to conserve the methylation information, Methyl-SNP-seq can be combined with DNA amplification using high throughput Illumina sequencer. Furthermore, the dual readout can be used both at the experimental and analytical steps. Experimentally, the original sequence can be used to enrich targeted regions using conventional 4-nucleotide probes matching the original sequence instead of bisulfite-converted probes that are specific to methylation. Subsequently, the methylation and variation can be phased together at molecule resolution while the redundancy of the complementary strands increase variant detection. In addition, Methylation motifs can also be directly detected from the sequencing reads without a reference genome.

**Figure 1.**
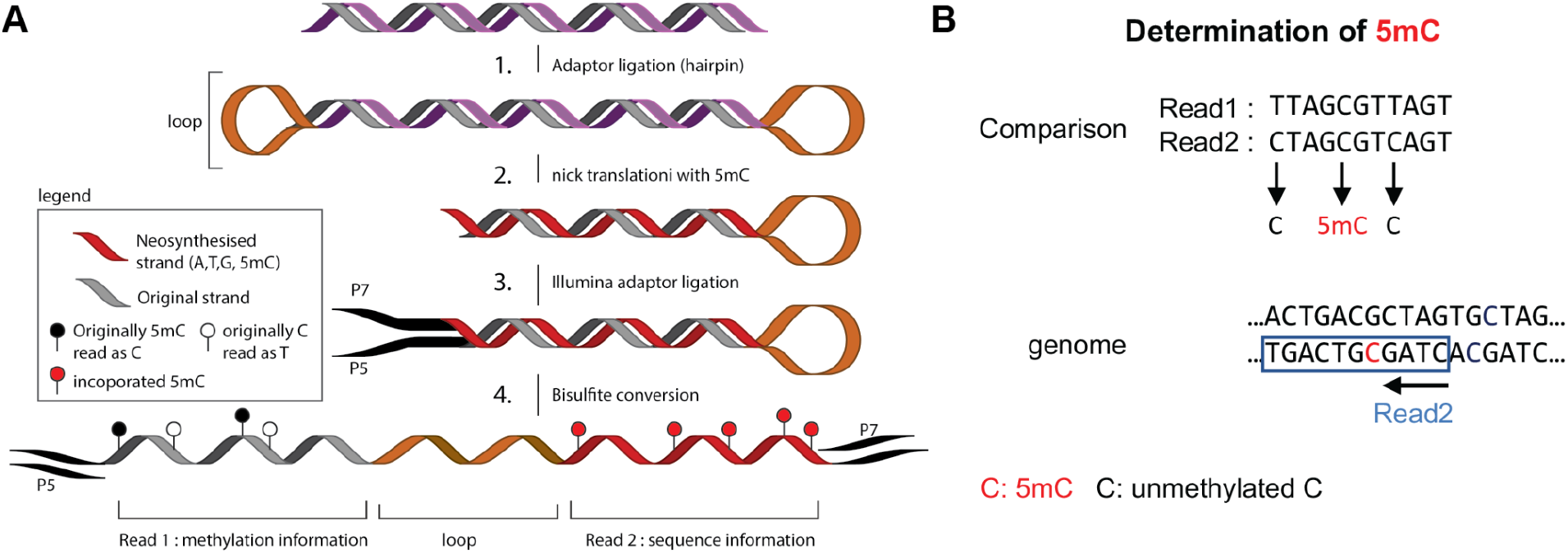
Overview of Methyl-SNP-seq : (**A)** Experimental workflow of Methyl-SNP-seq : 1-the genomic DNA is fragmented to 300-400bp-fragments. 2-Hairpin adapters are ligated at both ends of the fragmented DNA, forming a dumbbell shaped DNA. Next, nicks at both opposite ends of the adapters are introduced and using nick translation, a copy of the original strand is synthesized replacing CTP as a source of nucleotide with m5CTP instead. This nick translation step breaks the dumbbell shaped DNA somewhere within the fragment. 3-Methylated illumina Y-shaped adapters are ligated-. 4-bisulfite conversion opens the DNA structure revealing a single strand DNA molecule that can be amplified using the illumina adaptors. Sequencing requires paired-end reads to obtain both the methylation and the genomic sequence information. For more details on the experimental procedure, see Supplementary Figure 1A and Supplemental Protocol. **(B)** Deconvolution procedure: for more details on the bioinformatics analysis, see Supplementary Figure 1B.

We demonstrated the applicability of Methyl-SNP-seq for human genome sequencing with readouts that are comparable to commonly used techniques for both SNPs and methylation analysis. We also took advantage of the dual readout to accurately call Allele specific methylation (ASM). To demonstrate the efficient capture using conventional 4-nucleotide probe sets, we performed exome sequencing of a Methyl-SNP-seq library resulting in enrichment metrics that are comparable to standard DNA-seq based exome sequencing. Finally, we applied Methyl-SNP-seq on a bacterium and a synthetic microbial community and demonstrated the identification of methylation sites and methyltransferase specificity directly on reads without the need of a reference genome or assembly.

## Result

### 1. Principle of Methyl-SNP-seq

To capture the original four nucleotide sequence information as well as cytosine modification, Methyl-SNP-seq is taking advantage of the double stranded nature of DNA to duplicate the sequence information into a linked copy of the original strand. Importantly, this copy is amended to be resistant to bisulfite conversion by replacing all the cytosines with deamination-resistant modified nucleotides. Thus, the copied strand conserves its original four nucleotide content while the original strand undergoes deamination at unmethylated cytosines. Both strands are connected as one molecule and are sequenced using Illumina paired-end sequencing resulting in one read containing the sequence information while the other paired-read contains the methylation information (**Figure 1A and Supplemental Figure 1A**).

To achieve this, a hairpin adapter is ligated to both ends of the fragmented double stranded DNA, forming a dumbbell shaped DNA. Next, a nick at both opposite ends of the adapters is introduced, and a copy of the original strand is synthesized via nick translation while the other strand remains unchanged. Nick translation is performed in the presence of 5mdCTP as substrate and thus, the newly synthesized strand is resistant to bisulfite conversion. The nick translation step breaks the dumbbell shaped DNA somewhere within the fragment, creating two molecules each containing a neo-synthesized strand and the original strand. Methylated illumina Y-shaped adapters are ligated before bisulfite conversion. Bisulfite conversion deaminates unmodified cytosines to uracils on the original strand and opens the closed DNA structure, revealing a single strand DNA molecule that can be amplified using the illumina adaptors. Sequencing requires paired-end reads to obtain both the methylation and the genomic sequence information (**Methods**). We designed the protocol so that the Read1 of the paired-end read pair provides the cytosine methylation information conveyed by bisulfite conversion while the corresponding Read2 provides the genome sequence. To combine both information together, we developed a deconvolution algorithm (**Methods, Figure 1B and Supplemental Figure 1B**) that compares Read1 with Read2 considering the conversion and complementary nature of the paired-end reads. This step, called the read deconvolution step, accurately identifies each cytosine and its methylation status. More specifically, a T in Read1 pairing with a C in Read2 corresponds to an unmethylated C, while a C in Read1 pairing with a C in Read2 corresponds to a methylated C (**Figure 1B**) while all remaining pairs should match.

A typical Methyl-SNP-seq experiment yields about 85-90% of the reads being deconvoluted. Within the deconvoluted reads, around 98-99% of the positions show either a direct agreement between pairs or a profile consistent with cytosine conversion. The remaining 1-2% of bases that disagreed may be resulting from damages caused by the bisulfite reaction or errors generated during nick translation, PCR amplification or sequencing. In this case, we cannot differentiate the correct base. Accordingly, we use the Read1 base as the deconvoluted base but adjust the Phred quality score to mark this disagreement as a potential error. Adjustment of the Phred quality scores in the case of a pair disagreement depends on whether a reference genome is available or not. If a reference genome is available (Reference-dependent Read Deconvolution), base calibration is performed using Bayesian statistics which considers the corresponding nucleotide on the reference genome, the substitution type and sequencing cycle. Thus, the adjusted Phred quality score reflects the Bayesian probability that the base recorded on Read1 is true. If a reference genome is unavailable (Reference-free Read Deconvolution), the Phred quality score, in case of pair disagreement, is assigned to 0.

The deconvolution step results in a fastq file that contains deconvoluted reads with adjusted Phred quality scores and, for each cytosine, its methylation status in a methylation report file (**Supplemental Figure 1**). The pipeline for processing and deconvoluting the linked paired-end reads is freely available in Github (**Methods**). The outputs of the deconvolution pipeline are in a standard format compatible with existing algorithms designed for genome assembly, genetic variant calling (e.g. GATK (McKenna et al. 2010)) and methylation quantification (e.g. Bismark (Krueger and Andrews 2011)). The ability to distinguish between a methylated and unmethylated cytosine directly on the unmapped read while simultaneously obtaining the original genomic sequences is the key strength of this technology.

### 2. Application of Methyl-SNP-seq to whole genome sequencing of human GM12878 genomic DNA

As proof of concept, we tested Methyl-SNP-seq using gDNA from the widely studied human cell line GM12878 (lymphoblastoid cell line) for which a large number of sequencing and methylation datasets are publicly available. Unmethylated lambda DNA was spiked into the human gDNA to monitor the bisulfite conversion efficiency. Methyl-SNP-seq libraries were performed in duplicate using the same source of starting material to monitor the reproducibility of the method (**Methods**). Whole genome sequencing was done using Illumina Nova-seq resulting in an average of 1.5 billion 100bp paired-end reads per replicate. During the deconvolution step of Methyl-SNP-seq, an average of 84% of reads were successfully deconvoluted and more than 95% of the deconvoluted reads were mapped to the reference human genome using bowtie2 (Langmead and Salzberg 2012). To obtain a set of high confidence genetic variants and accurate methylation quantification, we applied stringent data filters to remove multiple mapping reads, PCR duplicates and reads indicating incomplete bisulfite conversion. About 64% of mapped reads remained after applying these filters. Supplemental Figure 1B shows the data analysis workflow used for this experiment. Both replicate show similar QC and alignment metrics (**Supplemental Table 1**) indicating that the Methyl-SNP-seq protocol is reproducible.

#### 2.1 Methyl-SNP-seq accurately detects genetic variation

We first assessed the ability of Methyl-SNP-seq to detect genetic variations in the human GM12878 cell line. To increase coverage, filtered reads from the two replicates were combined for variant calling. Genetic variants were identified using the GATK pipeline (McKenna et al. 2010) following the recommended best practice workflow. The resulting variants were benchmarked against the variants obtained using the NA12878 whole genome sequencing dataset (WGS, performed by JIMB NIST project). The number of true positive, false positive and false negative variants found using Methyl-SNP-seq were derived from the comparison between the two datasets. We first confirmed that the reference-dependent read deconvolution increases the number of true positive SNPs and reduces the number of false positive SNPs compared to reference-free read deconvolution (**Supplemental Figure 2A**).

Since both Deconvoluted Read and Read2 represent the original genome sequence, we can call genetic variants using either one. Overall, variants found using either the Deconvoluted Read or Read2 show a high level of agreement: 95% of SNPs found with the deconvoluted Read being identical to those found with Read 2 (**Figure 2A**). According to the experimental design, as expected the Deconvoluted Read and Read2 had different types of false positive errors (**Supplemental Figure 2B**). Consequently, by using the common set of variants defined by both deconvoluted read and Read2, we could correct the variant calling error and improve the accuracy (**Supplemental Figure 2C**). Therefore, we chose the common variants between Deconvoluted read and Read2 as the Methyl-SNP-seq defined genetic variants. Using this set of common variants, we found 1,296,699 and 1,903,083 homozygous and heterozygous SNPs respectively (vcf file available in GEO under accession number GSE206253). A total of 98% of the SNPs identified using Methyl-SNP-seq were confirmed by JIMB WGS with a better agreement for homozygous SNPs (accounting for a fifth of the total false positive SNPs) compared to heterozygous SNPs (accounting for four fifth of the total false positive SNPs) (**Figure 2A, C**). Our method also shows high accuracy for indels, with 94% agreement with WGS. These levels of agreement are comparable to those typically observed between standard WGS (Zook et al. 2014).

**Figure 2.**
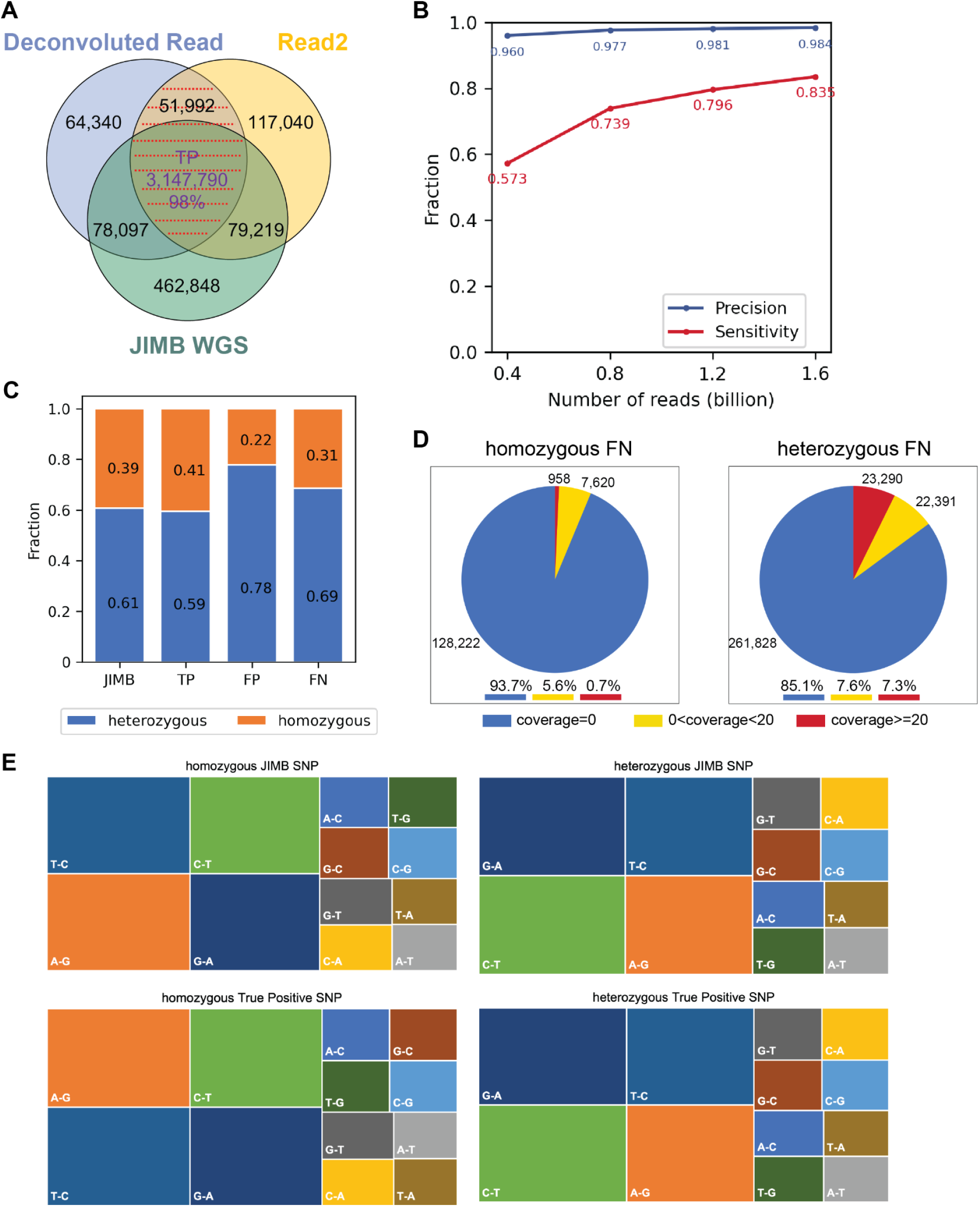
SNP identification by Methyl-SNP-seq. The JIMB whole genome sequencing of NA12878 was used as a benchmark for comparison. (**A)** Comparison of SNPs identified using Methyl-SNP-seq Deconvoluted Read and Read2 with those using JIMB whole genome sequencing data. Common SNPs, which were identified by both Deconvoluted Read and Read2 and marked by red dashed lines, are referred to as Methyl-SNP-seq defined SNPs. **(B)** Precision and Sensitivity of SNP identification using different numbers of Methyl-SNP-Seq reads. Precision=TP/(TP+FP). Sensitivity=TP/(TP+FN) with TP: True positive. FP: False positive. FN: False negative. **(C)** Fraction of heterozygous and homozygous Methyl-SNP-seq defined SNPs. **(D)** Distribution of the genome coverage of the False Negative SNP sites. **(E)** Characterization of the JIMB and True Positive Methyl-SNP-seq defined SNPs. (i.e. T-C means in the vcf file REF=T while ALT=C).

We performed standard quality controls for variant calling (Picard CollectVariantCallingMetrics) and found that both the Methyl-SNP-seq and the WGS datasets displayed comparable metrics for both SNPs and indels (**Supplemental Table 2**). More specifically, the ratio of transition (Ti) to transversion (Tv) mutations is around 2.06 for both datasets demonstrating that both sets are unlikely to have a bias affecting the transition transversion ratio. As expected, we have less accuracy in detecting the C-T (REF-ALT in vcf), T-C, G-A and A-G type SNPs, which, combined, account for most of the errors (**Supplemental Figure 3**).

To assess the sensitivity of Methyl-SNP-seq in identifying variants, we downsampled the dataset to various fractions of total reads. At equivalent coverage, we detected more than 80% of the WGS SNPs. This number drops to 60% when using only 25% of reads (**Figure 2B**). We noted that the lack of read coverage was the major cause of false negative SNPs from Methyl-SNP-seq (**Figure 2D**). Although having the same number of reads, the number of total bases was fewer in deconvoluted reads (109 billion) compared to WGS (160 billion) due to the shorter read length after trimming the hairpin adapter. Importantly, reducing the number of reads did not affect the accuracy for variant detection (**Figure 2B**).

#### 2.2 Methyl-SNP-seq accurately detects and quantifies cytosine methylation at base resolution

We next evaluated the performance of Methyl-SNP-seq in identifying and quantifying cytosine methylation. The methylation status of individual cytosine was determined in the Read Deconvolution step and was added in the mapped bam file so that the base-resolution methylation information can be calculated using the conventional methylation calling tools such as bismark (Krueger and Andrews 2011) (**Methods**). We benchmarked our method’s performance against two reference datasets generated by standard whole genome bisulfite sequencing method (ENCODE Project Consortium 2012) (Davis et al. 2018) and Nanopore sequencing (Jain et al. 2018). Using the unmethylated Lambda spiked-in control we estimated the bisulfite conversion rate of Methyl-SNP-seq to be 97.5%. Overall CpG methylation is at 45.3% for both replicates in line with the two ENCODE datasets analyzed showing 45.5% and 50.1% overall CpG methylation respectively. The GC bias of Methyl-SNP-seq follows closely the known GC bias observed for bisulfite sequencing (Olova et al. 2018) with a preferential sequencing of AT rich genomic regions. Both replicates show comparable results (**Supplemental Table 3**).

With 1.6 billion reads from the two replicates combined, we acquired 54 million CpG sites, for which 45 million had at least 5X coverage (methylation file available in GEO under accession number GSE206253). These numbers are comparable with that of the WGBS method down-sampled to the similar number of reads (53 millions sites with >=5X coverage) (**Figure 3A, B**). The genome-wide methylation level of CpG sites identified by Methyl-SNP-seq displays a bimodal distribution similar to those of the WGBS and Nanopore datasets (**Figure 3C**) with a distribution that better resembles the Nanopore dataset. This result agrees with the observation that bisulfite sequencing overestimates the global methylation (Jain et al. 2018; Olova et al. 2018) (Ji et al. 2014).

**Figure 3:**
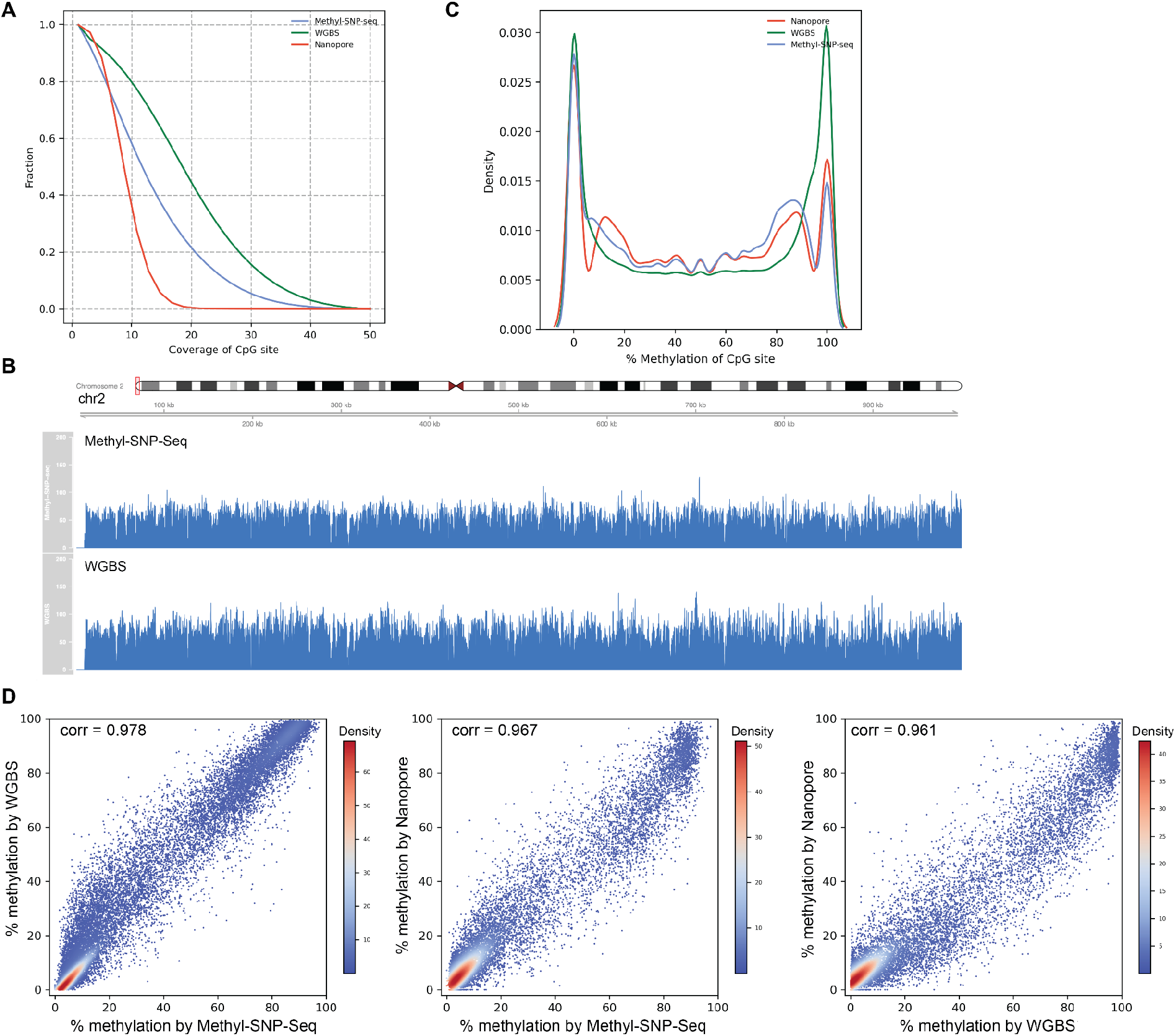
Methylome. **(A)** Distribution (kde plot) of % methylation on CpG sites having coverage>=5. **(B)** The genome coverage of Methyl-SNP-seq and WGBS on chr2. **(C)** Fraction of coverage on CpG sites. **(D)** Pairwise comparison of methylation level of CpG islands measured by Methyl-SNP-seq, whole genome bisulfite sequencing from ENCODE (WGBS) and Nanopore sequencing. Each dot represents the percentage methylation at CpG island. Only CpG islands having coverage>=50 were used for correlation calculation. A total of 27,050; 27,313 and 16,071 CpG islands were detected by Methyl-SNP-seq, WGBS and Nanopore sequencing respectively.

Methylation patterns of CpG islands have been shown to affect gene expression and are linked to disease phenotypes (Robertson 2005). Therefore, we compared the methylation level of the known CpG islands obtained using the three methods. Methyl-SNP-seq is highly correlated with both the ENCODE WGBS (Pearson correlation= 0.98) and Nanopore (Pearson correlation= 0.97) datasets (**Figure 3D**), indicating that Methyl-SNP-seq is highly accurate for cytosine methylation quantification.

#### 2.3 Allele-specific methylation using Methyl-SNP-seq

Attempts to infer SNPs from WGBS have been previously published (Liu et al. 2012a) but require high genome coverage (e.g. >30X coverage required by Bis-SNP) to assess independently paired-end reads. This is because the identification of transition SNP such as C/T, G/A, A/G and T/C are confounded by the deamination step (Liu et al. 2012b). In contrast, our method can confidently distinguish cytosine methylation from original transition SNPs along with other SNP types. Indeed, by using the redundancy of the double stranded DNA to read methylation and sequence from the same original DNA molecule, our method identifies both the methylation state and variants at single molecule level. This allows phasing of the methylation state with heterogeneous SNPs directly on the read, enabling the identification of differentially methylated genomic regions (DMR) that are allele specific (ASDMR).

Using the whole genome Methyl-SNP-seq experiment done on human GM12878 described above, we identified a total of 34,909 ASDMR genomewide. **Figure 4A** shows an example of a known ASDMR (Suzuki et al. 2018; Kaplow et al. 2015) containing the heterozygous SNP rs11686156 on chromosome 2. Among all the identified ASDMRs, 47% have SNP directly affecting CpG sites. This result is consistent with a previous study (Shoemaker et al. 2010), which reported that 38% to 88% of ASM regions are solely due to the presence of SNPs at CpG dinucleotides and indicates variation at CpG sites is a dominating factor for ASDMR. In this case, SNP not only disrupts the methylation pattern of the affected CpG site but also affects the methylation pattern of the neighboring regions. Therefore, CpG-SNPs are very important for DMR studies because they may play a role in the establishment of certain types of DMRs such as ASDMRs.

**Figure 4.**
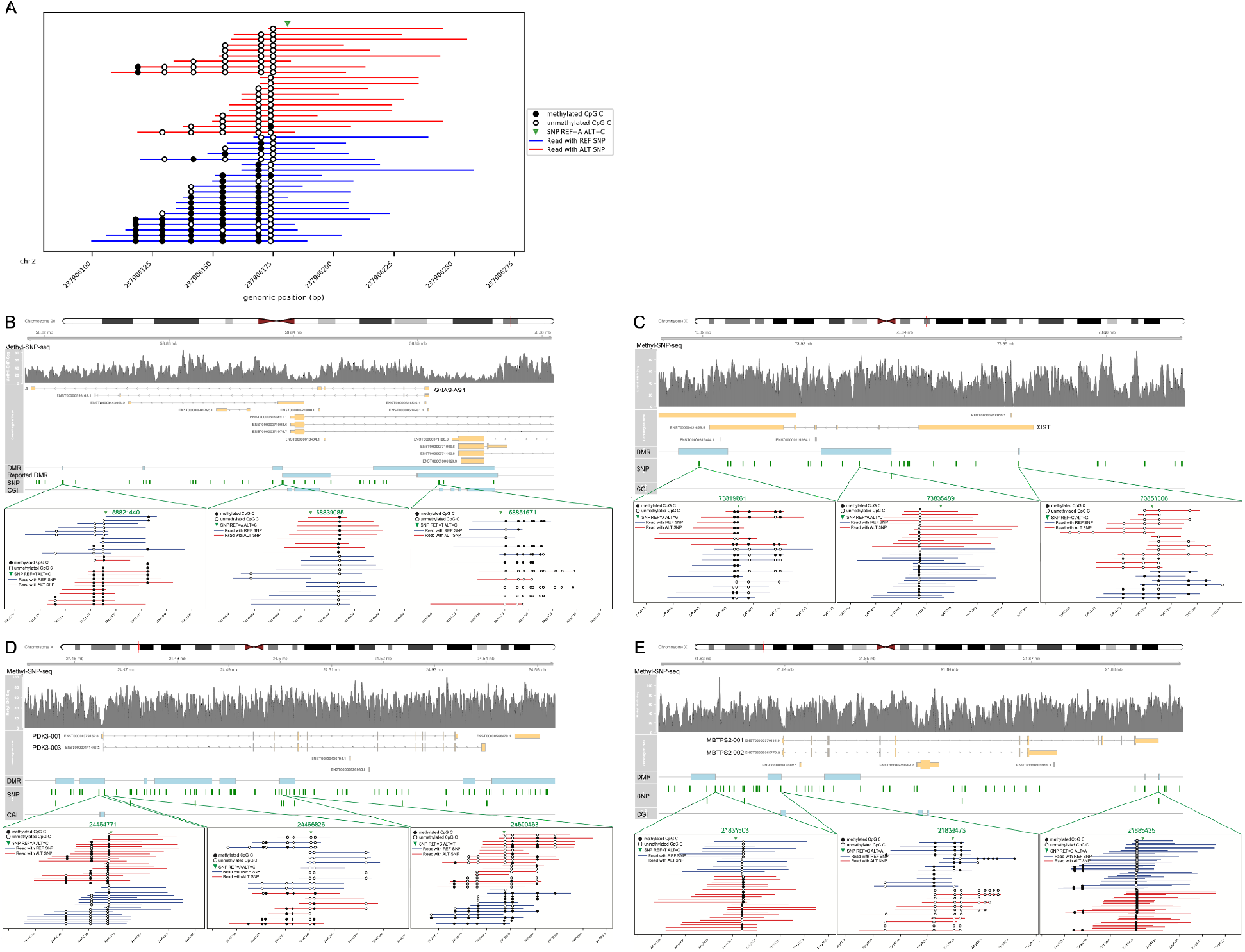
DMR. **(A)** chr2: A known example of allele-specific methylation. The reads having different alleles of the arrow pointed heterozygous SNP site are labeled with blue (REF SNP) and red (ALT SNP). SNP: rs11686156. **(B)** chr20: 10kb upstream of GNAS-AS1 gene and overlapping reported DMRs (Fang et al. 2012). **(C)** chrX: XIST. **(D)** chrX: PDK3. **(E)** MBTPS2. Only C in CpG contexts are shown. Coordinates on the x axis correspond to positions on respective chromosomes (GRCh38 assembly).

Allele-specific methylation is also often associated with gene imprinting. Using a set of ASDMRs that are reported to be associated with known imprinted gene clusters in the human genome as a reference (Fang et al. 2012), we were able to identify 24 ASDMRs at or near the reported imprinting control DMRs for 15 out of the 30 imprinted gene clusters (**Supplemental Table 4**). For example, we detected 2 ASDMRs overlapping with the known imprinted cluster of the GNAS gene (**Figure 4B**). These two ASDMRs span a 17.8kb region and include 670 CpG pairs. When examining the methylation level of all the CpGs within this region, we saw that most of the CpG sites have an average methylation level close to 50% whereas CpG sites in the flanking regions have elevated methylation levels, suggesting this entire 17.8kb region is likely an imprinted DMR (**Supplemental Figure 4**).

Allele-specific methylation (ASM) is also known to be associated with X chromosome inactivation in female cells via regulating the X-inactive specific transcript (XIST) gene (Wutz 2011; Fang et al. 2012). Accordingly, our method detected several ASM near the XIST gene in the human lymphocyte cell GM12878 (female) (**Figure 4C**). In addition, we also detected ASMs in the promoter regions of genes that are known to be subject to X-chromosome inactivation (XCI) (Cotton et al. 2015)(Sharp et al. 2011) such as PDK3 and MBTPS2 (**Figure 4D, E**)

A previous study (Kaplow et al. 2015) found that genomic regions with chromatin states consistent with active transcription and active enhancers were enriched for CpGs with mQTLs (ASM), suggesting that some of these ASM may affect transcription or enhancer activity. In our study, we found that CpGs that are associated with ASDMRs are significantly enriched, compared to random CpG regions, in enhancers which include both active and primed enhancers and are marked by histone H3K4me1 modification in the absence of histone H3K4me3 modification (χ2 =98.3, df=1, P-value < 1e-9, fold change=1.5) (**Supplemental Table 5 in Supplemental Material**). However, ASDMR CpGs are not enriched in active enhancers identified by H3K27Ac modification (fold change=0.9). In addition, ASDMR CpGs are significantly depleted in the promoter regions marked with histone H3K4me3 modification (χ2 =120.1, df=1, P-value < 1e-9, fold change=0.7). Interestingly, ASDMR CpGs are also enriched in the genomic regions with repressive histone mark H3K9me3 (χ2=29.1, df=1, P-value = 6.8E-8, fold change=1.4). This histone mark is associated with heterochromatin and frequently coexists with DNA methylation. H3K9me3 is also reported to play a role in establishing imprinted X-chromosome inactivation in mice (Fukuda et al. 2014).

### 3. Methyl-SNP-seq can be performed in conjunction with the conventional probe-based target enrichment

While providing a comprehensive view of the human genome, whole-genome sequencing remains cost-prohibitive for analyzing a large number of clinical samples. In contrast, targeted sequencing with a focus on specific regions of interest is more widely and commonly used. In particular, targeted bisulfite sequencing is designed to measure site-specific DNA methylation changes. Accordingly, it normally requires the design of specific bait probes capturing the bisulfite converted regions. Unlike the conventional targeted bisulfite sequencing, Methyl-SNP-seq method contains the original genome sequence (**Figure 1**) that can hybridize with the standard bait probes. Thus, in theory, Methyl-SNP-seq can be easily adapted to the conventional targeted enrichment method with any standard probe set.

To demonstrate the applicability of Methyl-SNP-seq for target enrichment, we tested Methyl-SNP-seq combined with the Twist human comprehensive exome panel. The targeted Methyl-SNP-seq had a high mapping efficiency with 96% of the reads mapping to the human genome. Bisulfite conversion rate is 97% at CHG and CHH contexts. Targeted-Methyl-SNP-seq also showed comparable target enrichment capability (**Supplemental Table 6**) compared to the standard exome targeted capture with the exception that it had lower coverage of AT-rich regions (AT_DROPOUT=9). We also applied stringent filters to remove PCR duplicates etc, consequently having about 11 million deconvoluted reads from two replicates. Although the Twist exome panel is designed to capture the gene body rather than the promoter regions, we still found 11783 CpG islands captured with coverage above 50. Like the whole genome sequencing, the methylation quantification of these CpG islands was consistent with the WGBS and Nanopore result (**Figure 5A**). As for the variant detection, using the same probe panel at equivalent read depth (about 10 million reads used for variant calling), the precision of targeted Methyl-SNP-seq sequencing (precision = 0.8) was lower than the standard targeted sequencing (precision = 0.9) (**Figure 5B**) (Zhou et al. 2021).

**Figure 5.**
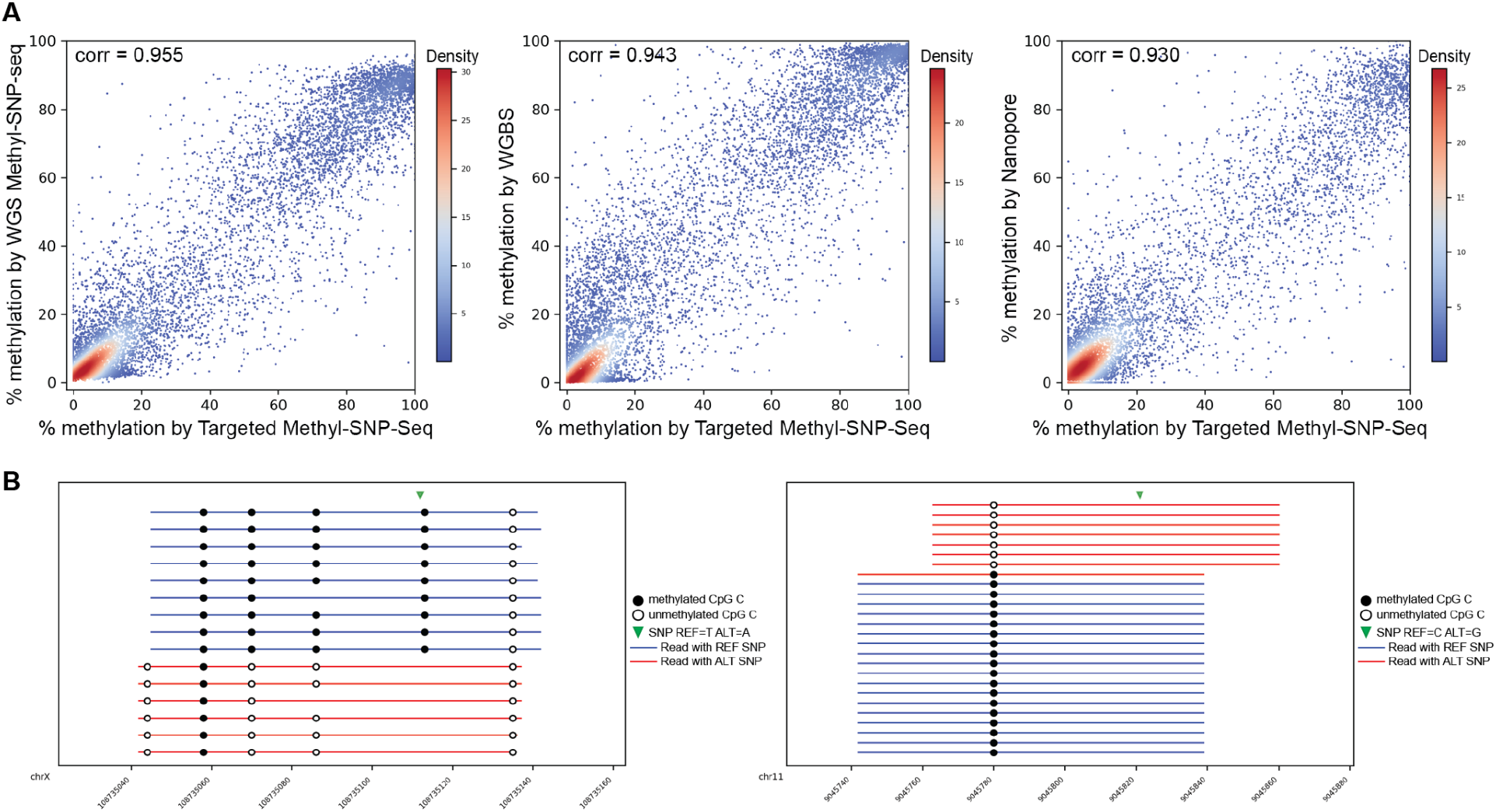
**(A)** Pairwise comparison of methylation level of CpG islands measured by targeted Methyl-SNP-seq, whole-genome Methyl-SNP-seq (WGS Methyl-SNP-seq), whole-genome bisulfite sequencing by ENCODE (WGBS) and Nanopore sequencing. Only CpG islands having coverage>=50 were used for correlation calculation. **(B)** Two ASM depicted by targeted Methyl-SNP-seq. Only C in CpG contexts are shown. Coordinates on the x axis correspond to positions on X (felt) and chr11 (right) (GRCh38).

### 4. Reference-free identification of m5C in bacteria using Methyl-SNP-seq

Another application of Methyl-SNP-seq is on the identification of methylation in organisms for which a reference genome or assembly is missing. This is often the case for environmental samples and microbiomes. In these cases, conversion-based methods to identify methylation (e.g. bisulfite sequencing) cannot be used because these methods rely on differentiating between a genuine T and a C to T conversions using a reference genome. The Methyl-SNP-seq method, on the other hand, identifies cytosine methylation directly on the paired-end reads in a reference-independent manner. Additionally, it reports methylation status of individual cytosine sites with sequence context information at single-base resolution and at single-molecule level, which is most suitable for methylation motif studies. Furthermore, our Methyl-SNP-seq method also reports the original genomic sequences that can be used for genome assemblies of a single organism or a mixed population.

To demonstrate the effectiveness of Methyl-SNP-seq for these applications, we performed Methyl-SNP-seq using genomic DNA of an isolated strain of *E. coli* K12. We first investigated whether we could assemble the deconvoluted reads into a reliable reference genome. Using the Velvet assembler (Zerbino 2010), we obtained a good assembly from the *E. coli* data (16 million deconvoluted reads) with high genome coverage (94% of the genome covered) and high sequence identity (2.21 mismatches per 100 kbp) (**Figure 6A**), which was comparable to the performance of the assembler for single-end short read assembly using standard DNA-seq.

**Figure 6.**
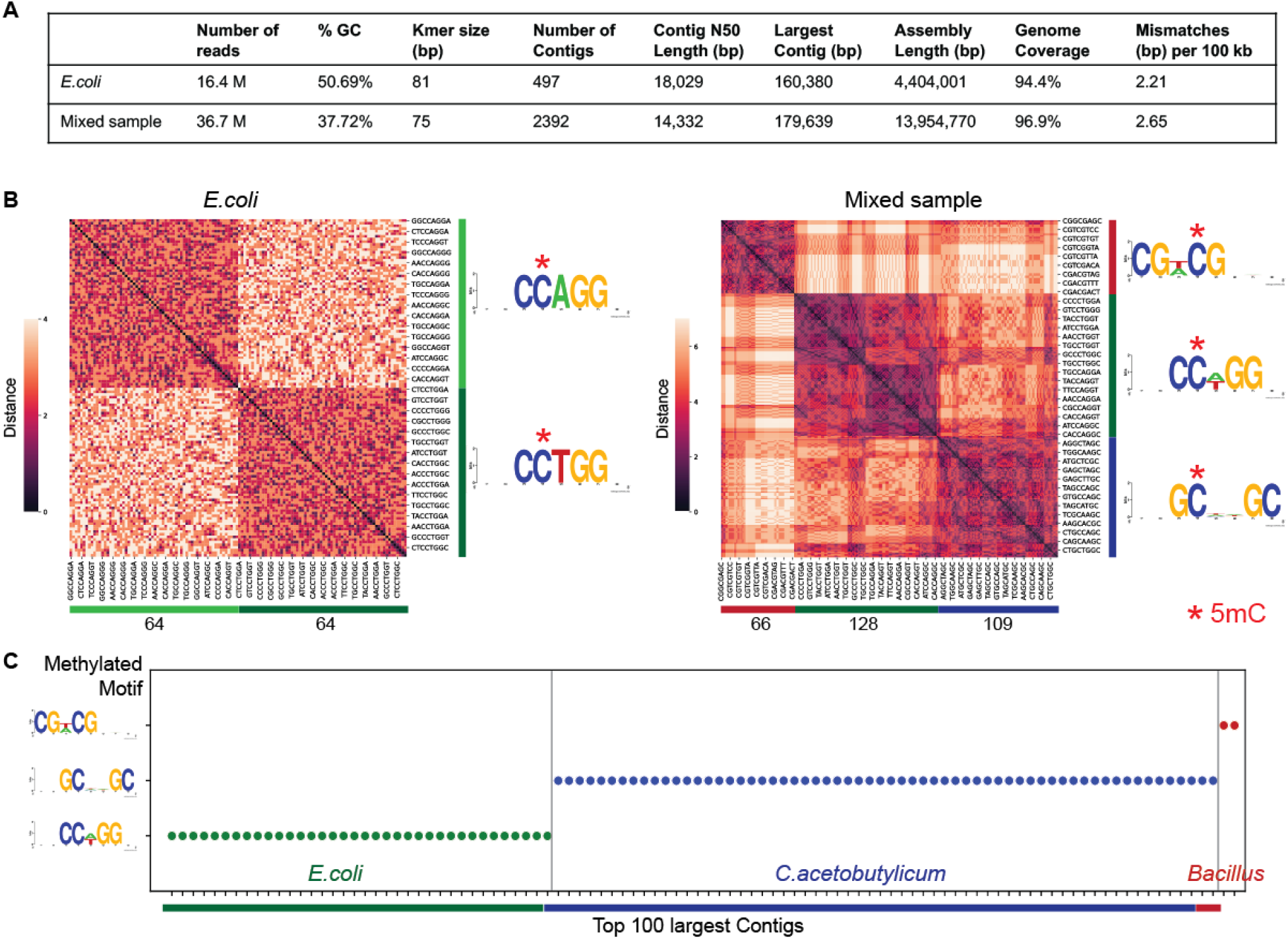
**(A)** Summary of genome assembly using Deconvoluted reads. Genome coverage was measured against *E. coli* or *E. coli* + *C. acetobutylicum* reference genome using Quast. **(B)** Hierarchy clustering of significantly enriched 8mers and the corresponding motif logo (generated by weblogo) for *E. coli* and mixed sample (*E. coli* + *C*.*acetobutylicum*). The number at the bottom represents the number of enriched kmers in each cluster. **(C)** Identified methyltransferase motif for the top 100 largest assembled contigs. Each dot represents a contig.

We next determined the methyltransferase specificity of the bacteria directly on the deconvoluted reads without mapping to a reference genome. To achieve this, we randomly selected 0.3 million deconvoluted reads and counted the number of occurrences of all the 8bp kmers (8mers) having methylated or unmethylated cytosine from these reads. We applied a Binomial model with Bonferroni adjustments to identify the 8bp sequences having significantly higher methylation levels. These sequences were further grouped using Hierarchical clustering to uncover the consensus methylated motif indicative of methyltransferase specificity(ies). Two context clusters were found from 128 significantly enriched 8mers. CCAGG and CCTGG which can be further combined into CCWGG (with W=A or T), revealing the correct specificity for the *E. coli* dcm methyltransferase (Marinus and Morris 1973; May and Hattman 1975) (**Figure 6B**).

We also performed Methyl-SNP-seq on a mixed sample consisting of genomic DNA of two bacterial strains (*E. coli* K12 and *C. acetobutylicum* ABKn8 strains) to mimic a simple mixed bacteria population. Using 0.6 million deconvoluted reads we found 3 motifs instead of the 2 motifs expected for these strains (**Figure 6B**). These motifs correspond to CCWGG, GCNNGC, and CGWCG. The first two motifs match the expected methyltransferase specificities of *E. coli* dcm and *C. acetobutylicum* (Baum et al. 2021), respectively. The third motif, CGWCG, however, was unexpected. To investigate the origin of this methylated motif, we assembled the deconvoluted reads of the mixed sample library and used the 100 longest assembled contigs as reference to determine methylation motifs. All of the 100 contigs contain a single methylated motif, among which 36 and 62 have the CCWGG motif and the GCNNGC motif, consistent with their *E. coli* and *C*.*acetobutylicum* origins (**Figure 6C**). Two out of the 100 contigs have the unexpected CGWCG motif and both contigs have high sequence homology to the genome sequence of a *Bacillus* strain by BLAST search, implying that there was a contamination and most likely a *Bacillus* strain in the mixed sample. We further annotated the assembled contigs which contain the CGWCG motif using Prokka and identified a single gene with cytosine-specific DNA methyltransferase domain. This suggests the contamination strain is methylated. Additionally, the methylation analysis using EM-seq confirmed the presence of a CGWCG methylation motif in the used *C*.*acetobutylicum* sample.

In this example, we demonstrated that Methyl-SNP-seq method can identify all the methylation motifs from a mixed sample in a reference-independent manner, as well as resolve the composition of a mixed population by assembling the deconvoluted sequences and using methylation motif as a species/strain signature and genome binning criteria.

## Discussion

Amplification-based sequencing methods provide only the sequential arrangement of the canonical four bases A, T, C and G while all modifications, originally present on the DNA, are erased. The information on what base was originally modified is lost during the in-vitro DNA synthesis steps that happen during amplification, clustering, and sequencing. To circumvent this limitation and obtain cytosine methylation information, techniques such as bisulfite sequencing convert unmethylated Cs to Ts before subjecting the converted DNA to sequencing. A T output after bisulfite treatment is therefore ambiguous: it corresponds to either a naturally occurring T in the sequence or a deaminated unmodified C and a reference genome is therefore required to distinguish the two possibilities. This ambiguity is the major drawback in bisulfite sequencing and relegates all the techniques that rely on deamination to applications directed for methylation analysis only.

By locking the double strands together, Methyl-SNP-seq takes advantage of the redundant information captured in the complementary strands to obtain both the arrangement of the canonical four bases and the methylation information. The accuracy of the dual readouts of Methyl-SNP-seq is comparable to state-of-the-art techniques for both SNPs and methylation analysis. Because the sequencing power is allocated to a dual readout, the sensitivity for every single readout is reduced to effectively a single-end read instead of a paired-end read. This affects notably the ability to perform assemblies as most of the assemblers have been optimized for paired-end sequencing. With the ability to read longer stretches of sequence, this limitation can be partially overcome. Furthermore, there should be no technical limitations from the manufacturer in adapting the instrument to perform dual paired-end read sequencing using primer targeting the invariable hairpin structure. The dual readout also implies that the sequencing volume to achieve the similar sensitivity obtained by WGBS or DNA-seq needs to be increased.

Nonetheless, Methyl-SNP-seq is a single experiment that is more convenient than performing WGBS and DNA-seq separately. In addition, Methyl-SNP-seq offers important functionalities that are not feasible when performing WGBS or DNA-seq. Notably, Methyl-SNP-seq preserves the base arrangement of one of the double strands by incorporating m5CTP instead of CTP in the neo-synthesized fragment. This is conceptually a significant improvement compared to a previously published method (Liang et al. 2021) for which both strands are subjected to deamination. In the latter case, the ability to obtain the original sequence can only be done computationally, by aligning and deconvoluting paired-end reads. In addition, by keeping intact the 4-nucleotide-based strand, Methyl-SNP-seq is compatible with conventional probe sets for target enrichment. Indeed, we show similar on-target performance for both conventional DNA-seq and Methyl-SNP-seq exome sequencing. Methyl-SNP-seq exome sequencing shows a SNP detection accuracy lower than conventional exome sequencing. Thus, a higher coverage is likely needed to obtain similar SNP accuracy.

Retaining the original sequence is also useful for any target-specific amplification such as CRISPR-based targeting, and other sequence-specific technologies. Beyond sequence-specific applications, we demonstrate the applicability of Methyl-SNP-seq in directly demonstrating allele-specific methylation at single-molecule resolution. In conjunction with target-specific amplification and sequencing, Methyl-SNP-seq is an ideal technique to validate candidate ASMs derived from Methylome-Wide Association Studies.

Beyond human methylomes, Methyl-SNP-seq is a useful technology notably for organisms for which a reference genome is not available such as non-model organisms and microbial communities. Notably, the identification of modification directly on the unmapped reads enhanced the ability to bin sequences based on methylation patterns, an important feature for resolving genomes within a complex community (Wilbanks et al. 2022)(Tourancheau et al. 2021). The ability to obtain the original genomic sequence allows further functionalities specific to organisms for which a reference genome is unavailable or variations between the studied organism and its reference genome are too high to confidently distinguish methylation from transition SNPs. For example, we demonstrate the ability to perform assemblies and overlay methylation on the newly assembled genome.

## Methods

### Preparation of Methyl-SNP-seq library

For human Methyl-SNP-seq sequencing, we used genomic DNA isolated from the GM12878 cell line (NA12878, provided by Coriell Institute) for library preparation. For human whole-genome Methyl-SNP-seq sequencing, we used 4 ug of NA12878 gDNA and the unmethylated lambda DNA as spiked in to monitor the bisulfite conversion efficiency. The genomic DNA was fragmented using 250bp sonication protocol using a Covaris S2 sonicator. We set up two technical replicates. For human exome targeted Methyl-SNP-seq sequencing, 4 ug of NA12878 gDNA was fragmented using 400bp or 500bp sonication protocol.

For bacteria Methyl-SNP-seq sequencing, we used 2 ug of *E. coli* genomic DNA (MG1655 strain) or 2ug of mixed bacterial DNA (containing 1ug of *E. coli* MG1655 genomic DNA and 1ug of *C. acetobutylicum* genomic DNA). The genomic DNA was fragmented using 250bp sonication protocol.

As shown in **Supplemental Figure 1A**, the fragmented gDNA was end-repaired and dA-tailed (NEB Ultra II E7546 module), then ligated to a custom hairpin adapter using NEB ligase master mix (NEB, M0367). The incomplete ligation product (fragment having only one or no adaptor ligated) was removed using exonuclease (NEB ExoIII and NEB ExoVII). Two nick sites were created at the Uracil positions in the hairpin adapters at both ends after being treated with UDG (NEB) and endoVIII (NEB). The nick sites were translated towards 3’ terminus by DNA polymerase I in the presence of dATP, dGTP, dTGP and 5-methyl-dCTP. The nick translation causes double-stranded DNA break when DNA polymerase I encounter the other nick on the opposite strand. The resulting fragments have one end ligated to a hairpin adapter and a blunt end on the other side. The blunt end was dA-tailed and ligated with Y-shape methylated Illumina adapter. The ligated product was bisulfite converted using Abcam Fast Bisulfite conversion kit (Abcam, ab117127). The bisulfite converted product was amplified using NEBNext Q5U Master Mix (NEB, M0597). The resulting indexed library was used for Illumina sequencing or target enrichment.

To perform targeted sequencing, about 200-300 ng Methyl-SNP-seq indexed library was used in a pool for target enrichment. The whole human exome regions were enriched from the pooled libraries using the Twist Human Core Exome panel (Twist, 102025) following the manufacturer’s instructions. The enriched DNA fragments were further amplified using NEBNext Q5 Master Mix (NEB, M0544) and NEBNext Library Quant Primer Mix (NEB, E7603) for sequencing following the manufacturer’s instructions.

The human Methyl-SNP-seq libraries (WGS sequencing and targeted sequencing) were sequenced using an IlluminaIllumina Novaseq 6000 sequencer for 100bp paired-end reads. The bacteria Methyl-SNP-seq libraries (*E. coli* or mixed sample) were sequenced using an IlluminaIllumina Nextseq 550 sequencer for 150bp paired-end reads.

The sequence of the custom hairpin adapter (46bp) is:

5’-(p)CCACGACGACGACGACGAGCGTTAGGCTCGTCGTCGTCGTCGUGGT-3’

A detailed protocol is provided as Supplemental Protocol in Supplemental Material.

### Preparation of EM-seq library

We used 100 ng of *C. acetobutylicum* genomic DNA to prepare an EM-seq library (NEB E7120) as directed by the manufacturer. The library was sequenced using an Illumina Nextseq 550 sequencer for 75bp paired-end reads.

### Data analysis

- Reference genome and other annotation files GRCh38 human reference genome: ftp://ftp.ncbi.nlm.nih.gov/genomes/all/GCA/000/001/405/GCA_000001405.15_GRCh38/seqs_for_alignment_pipelines.ucsc_ids/GCA_000001405.15_GRCh38_no_alt_analysis_set.fna.gz Lambda reference genome: https://www.ncbi.nlm.nih.gov/nuccore/215104 Known human SNP files used for gatk Base Quality Recalibration: ftp://ftp.broadinstitute.org/bundle/hg38/dbsnp_138.hg38.vcf.gz ftp://ftp.broadinstitute.org/bundle/hg38/Mills_and_1000G_gold_standard.indels.hg38.vcf.gz ftp://ftp.broadinstitute.org/bundle/hg38/1000G_phase1.snps.high_confidence.hg38.vcf.gz JIMB NA12878 SNP vcf file used for ASM identification: ftp://ftp-trace.ncbi.nlm.nih.gov/giab/ftp/release/NA12878_HG001/NISTv3.3.2/GRCh38/HG001_GRCh38 _GIAB_highconf_CG-IllFB-IllGATKHC-Ion-10X-SOLID_CHROM1-X_v.3.3.2_highconf_PGandRTGphasetransf er.vcf.gz Human CpG island annotation was generated using the ‘CpG island track’ from UCSC browser.
- Data Processing for Methyl-SNP-seq The sequencing reads were trimmed for both Illumina adapter and hairpin adapter (https://github.com/elitaone/Methyl-SNP-seq/tree/main/Read_Processing/TrimRead.py) using Trimgalore version 0.6.4. For human NA12878 Methyl-SNP-seq sequencing, the bases of the last cycle [cycle 100] for both Read1 and Read2 were further trimmed due to poor quality. Next, we performed Read Deconvolution, which determines the base, adjusts the base quality scor,e and extracts the methylation information by comparing the paired Read1 and Read2. This step generates a fastq file containing the deconvoluted reads and a corresponding methylation report. The principle of Read Deconvolution is explained as follows (**Figure 1B**). Reference-free Read Deconvolution If the reference genome is not available, e.g. Methyl-SNP-seq for *E. coli* and mixed bacteria sample, a reference-free Deconvolution was performed using a custom pipeline (https://github.com/elitaone/Methyl-SNP-seq/tree/main/Read_Processing/DeconvolutionConversion_v2.py), including the following steps: (1) Base determination and methylation extraction. For the same Illumina cycle, if Read1 base is a C and Read2 base is a C, it results in a C in the deconvoluted read and a 5mC in the methylation report; while if Read1 base is a T and Read2 base is a C, it results in a C in the deconvoluted read and an unmethylated C in the methylation report. (2) Base quality score adjustment. For the mismatching positions that Read1 bases are different from Read2 bases except for the Read1-T Read2-C case, Read1 bases are used but the sequencing quality scores are adjusted to 0 in the deconvoluted reads. Reference-dependent Read Deconvolution (**Supplemental Figure 1B**) If the reference genome is available, e.g. Methyl-SNP-seq for human NA12878, a reference-dependent Deconvolution was performed to increase the performance (https://github.com/elitaone/Methyl-SNP-seq/tree/main/Read_Processing/DeconvolutionWithCalibration). The step (1) Base determination and methylation extraction is the same as the Reference-free Read Deconvolution. But Reference-dependent Read Deconvolution uses a statistical model for the base quality score adjustment in step 2 as shown below. (2) Base quality score adjustment. For the mismatching positions, by comparing to the reference genome, a Bayesian probability is calculated, which reflects the likelihood of being able to trust the Read1 base. Therefore, Read1 bases are used but the sequencing quality scores are adjusted based on the Bayesian probability in the deconvoluted reads.
- Alignment and Data Filtering for human NA12878 Methyl-SNP-seq (**Supplemental Figure 1B**) For human NA12878 Methyl-SNP-seq, the Deconvoluted Reads were aligned to the GRCh38 human reference genome using bowtie2 (version 2.3.0) default parameter for single end mapping with the addition of read group identifier defined by -- rg-id and --rg. These identifiers including the information for sequencing platform, flow cell and lane, barcode and sample were necessary for Base Quality Score Recalibration by gatk for variant calling. To achieve high accuracy, the following steps were taken to filter the aligned data before variant calling and methylation status determination:
  1. removal of multiple mapping using an in-house script (https://github.com/elitaone/Methyl-SNP-seq/tree/main/Read_Processing/MarkUniread.py). Here for bowtie2 single end mapping, the unique mapping is defined as the read having only AS tag or AS score != XS score (bowtie2 AS: best alignment score, XS: second-best alignment score);
  2. removal of PCR duplicates using an in-house script (https://github.com/elitaone/Methyl-SNP-seq/tree/main/Read_Processing/MarkDup.py). Here for bowtie2 single end mapping, the PCR duplicates are identified as reads aligned to the same position as well as having the same sequence;
  3. addition of XM tag reflecting the methylation status. Based on the Methylation report generated in the Read Deconvolution step, an XM tag is added to each mapped read in the sam file using an in-house script (https://github.com/elitaone/Methyl-SNP-seq/tree/main/Read_Processing/AddXMtag.py). The XM tag is defined by Bismark to mark methylation call string and used to extract methylation status;
  4. removal of reads having incomplete bisulfite conversion using Bismark (version 0.22.3) filter_non_conversion.
  The resulting filtered Deconvoluted Reads from two replicates were combined to be used for variant calling and methylation determination. There were 1.6 billion and 11 million filtered deconvoluted reads for human WGS and exome targeted Methyl-SNP-seq, respectively.
- Data Processing for human NA12878 whole genome sequencing Whole-genome sequencing of human NA12878 generated by JIMB NIST Genome in a Bottle (Zook et al. 2016) (JIMB WGS HG001) was used as a benchmark for comparison with Methyl-SNP-seq for variant calling. For a fair comparison to avoid differences due to the choice of variant calling pipeline (Cornish and Guda 2015), we processed the JIMB WGS data set using the same strategy as for the human Methyl-SNP-seq: (1) shortening the paired-end reads to 99bp; (2) trimming Illumina adapter; (3) bowtie2 mapping for the paired-end reads; (4) removing multiple alignments and PCR duplicates using samtools (version 1.14) markdup; (5) removing multiple mapping using the inhouse script (https://github.com/elitaone/Methyl-SNP-seq/tree/main/Read_Processing/MarkUniread.py). To achieve a similar coverage, we downsampled to use 1.6 billion filtered JIMB WGS reads for variant calling.
- Data Processing for human NA12878 whole-genome bisulfite sequencing Whole-genome bisulfite sequencing (WGBS) of human NA12878 generated by ENCODE (ENCSR890UQO) was compared with Methyl-SNP-seq for methylation quantification. For a fair comparison, we reduced the paired-end WGBS data to 99bp long and trimmed the Illumina adapters. Next, the read pairs were aligned to the human GRCh38 genome using Bismark (version 0.22.3). The properly paired reads were further filtered before methylation determination by: (1) removing PCR duplicates using samtools markdup; (2) filtering out alignments having incomplete bisulfite conversion using Bismark filter_non_conversion. The two ENCODE replicates were combined to about 1.6 billion filtered reads for methylation quantification.
- Variant calling and SNV comparison We performed variant calling on the filtered data set as mentioned above using gatk (version 4.1.8.1), following gatk best practice recommendations for germline short variant discovery. First, BaseCalibration (BaseRecalibrator and ApplyBQSR) was applied on the filtered data set to calibrate the systematic errors made by sequencing. Next, the calibrated reads were used for variant calling using HaplotypeCaller. Finally, FilterVariantTranches was applied to filter raw SNVs using --info-key CNN_1D and --snp-tranche 99 --indel-tranche 99. For human targeted Methyl-SNP-seq sequencing, an additional filter ‘DP < 6’ was applied to remove SNPs with low coverage. In this study, only SNVs on the somatic chromosomes, chrX and chrM were reported and used for analysis. The common SNVs identified by both Deconvoluted Read and Read2 were used as the Methyl-SNP-seq defined genetic variants. We used vcfeval from RTG Tools (version 3.11) (Cleary et al. 2014) to compare the SNVs defined by Methyl-SNP-seq or the benchmark JIMB WGS.
- Methylation quantification For Methyl-SNP-seq and WGBS, the methylation information was extracted on the filtered reads or read pairs using bismark_methylation_extractor (version 0.22.3) with the following parameters: --merge_non_CpG --bedGraph. We also used the latest Nanopore sequencing data set of the human GM12878 cell line for methylation detection (https://github.com/nanopore-wgs-consortium/NA12878/blob/master/Genome.md) (Jain et al. 2018). The Nanopore reads (in total 8.7 million from 21 runs) were aligned to the human GRCh38 genome using minimap2 (version 2.17). The methylation modification was detected using nanopolish (version 0.13.2) call-methylation function. The methylation level of UCSC annotated CpG islands (CGI) was defined as: CGI methylation = number of methylated CpG Cs in the region/number of CpG Cs in the region Only the CGIs having coverage (number of CpG Cs in the region) above 50 were used for comparison between different methods.
- Allele-specific methylation (ASM) determination To discover the allele-specific methylation loci in the NA12878 genome, we used the heterozygous SNPs detected by Methyl-SNP-seq and confirmed in the JIMB NA12878 SNP vcf file (Zook et al. 2019). We split the Methyl-SNP-seq alignments into two groups based on the defined SNP: REF (reads having the reference SNP) and ALT (reads having the alternative SNP). The methylation status of CpG sites was extracted for each group using bismark_methylation_extractor as previously mentioned. Finally, the differentially methylated regions between the REF and the ALT groups were detected using DSS tool (version 2.38.0) (Feng et al. 2014) with the following threshold for callDML and callDMR function: delta=0.1, p.threshold=0.05.
- Genome assembly of Methyl-SNP-seq of *E. coli* and mixed bacterial sample Bacterial genomes were assembled using velvet (version 1.2.10) based on 16.4 million and 36.7 million deconvoluted reads for *E. coli* and mixed samples, respectively. The following parameters were used for velvet assembly to obtain the best result: for *E. coli*, k=81 -fastq -short -exp_cov 13 -cov_cutoff 9 -min_contig_lgth 500; for mixed sample, k=75 -fastq -short -exp_cov 15 -cov_cutoff 8 -min_contig_lgth 500. The assembly quality was estimated using QUAST web interface (Gurevich et al. 2013).
- Determination of methyltransferase recognition site based on the deconvoluted reads We randomly chose 2% deconvoluted reads to identify the methyltransferase recognition site in *E. coli* or mixed samples (0.28 million or 0.67 million reads, respectively). The 8mers including 3bp upstream and 4bp downstream of either a 5mC or unmethylated C were extracted for each read. The reads including more than one methylated C were excluded from this analysis (https://github.com/elitaone/Methyl-SNP-seq/tree/main/Methylation_Motif_Calling/motifExtraction.py -l 3 -r 4). The numbers of 8mers containing either 5mC or 5mC and unmethylated cytosine were counted. We used Binomial statistics with Bonferroni Correction to determine the 8mer sequences that have significantly higher methylation level compared to the background. The p-value is calculated using the following formula. P-value (of each 8mer sequence) = 1 - binom.cdf(k, n, P0) For each 8mer sequence, k is the number of 8mers having 5mC; n is the number of 8mers having 5mC and unmethylated cytosine; P0 is the average methylation level. We used a custom script to perform this statistical analysis to extract the significantly enriched/methylated 8mers (https://github.com/elitaone/Methyl-SNP-seq/tree/main/Methylation_Motif_Calling/identifyMotify.py, --alpha 0.0001 for Binomial Significance Levels, --mode average). These significantly enriched 8mer sequences were further clustered to create the motif logo using a hierarchical linkage method based on the difference between each pair of sequences (https://github.com/elitaone/Methyl-SNP-seq/tree/main/Methylation_Motif_Calling/clusterMotif.py). The number of clusters (--number) can be decided based on cluster heatmap. Specifically, in this study we assigned the significantly enriched sequences into 2 clusters for *E. coli* and 3 clusters for the mixed sample (**Figure 6B**).

## Data Access

All raw and processed sequencing data generated in this study have been submitted to the NCBI Gene Expression Omnibus (GEO; https://www.ncbi.nlm.nih.gov/geo/) under accession number GSE206253.

## Conflict of interest statement

BY, RV, ZS, and LE are employees of New England Biolabs, Inc, a manufacturer of restriction enzymes and molecular biology reagents.

## Acknowledgments

We thank Alexey Fomenkov for providing the bacterial genomic DNA, Nan Dai and Ivan Correa for performing LC-MS on nucleotide pools, the NEB Sequencing core for performing Illumina sequencing and Chen Song for helpful suggestions on target enrichment strategies. This work was supported by New England Biolabs Inc.

## Notes

https://www.ncbi.nlm.nih.gov/geo/query/acc.cgi?acc=GSE206253

