## Supplemental Material for "Methyl-SNP-seq reveals dual readouts of methylome and variome at molecule resolution"

### Supplemental Figures

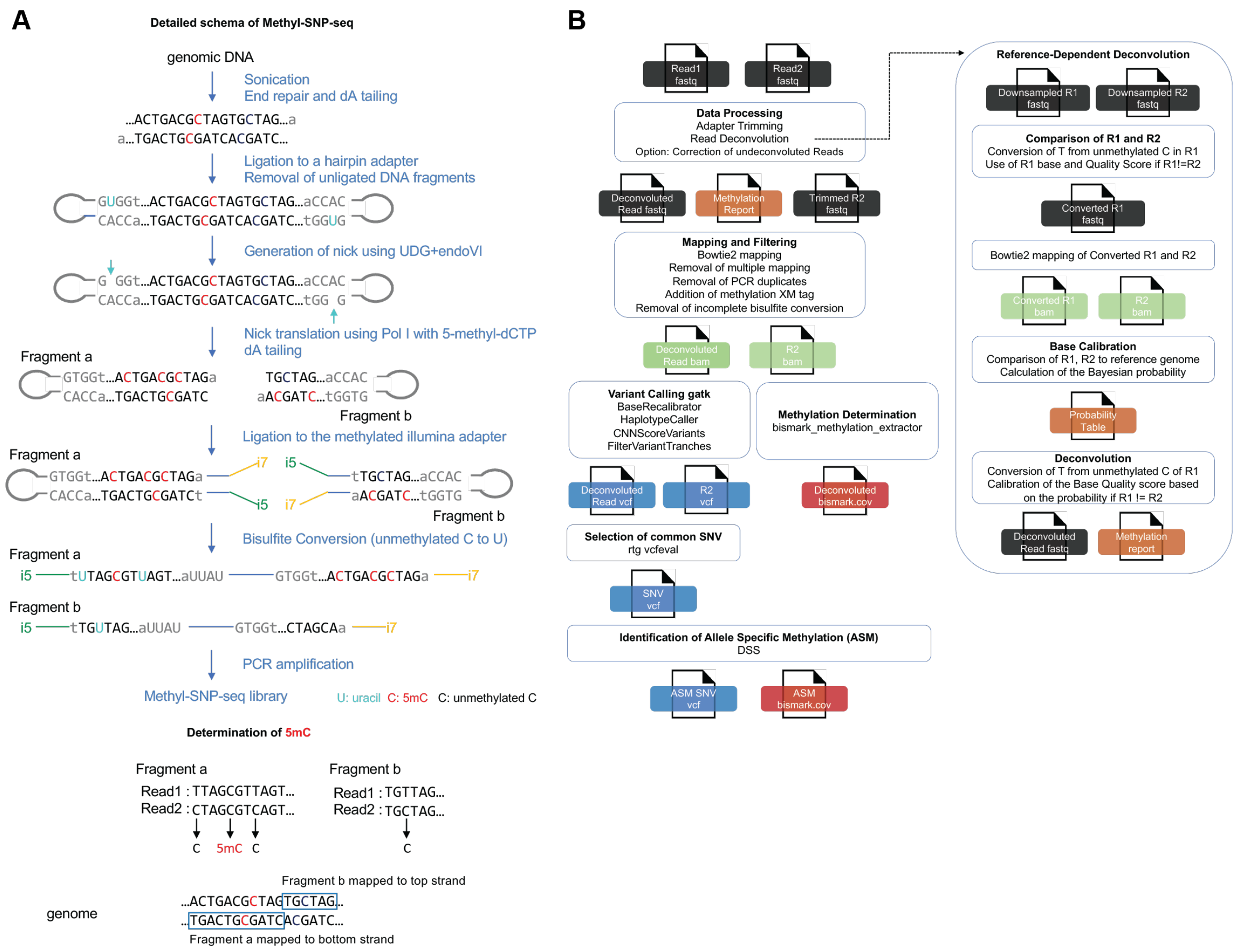


Supplemental Figure 1:
(A) Detailed description of the Methyl-SNP-seq experimental workflow. (B) Flowchart illustration of the analysis of Human Methyl-SNP-seq data. R1 and R2 stand for Read1 and Read2.

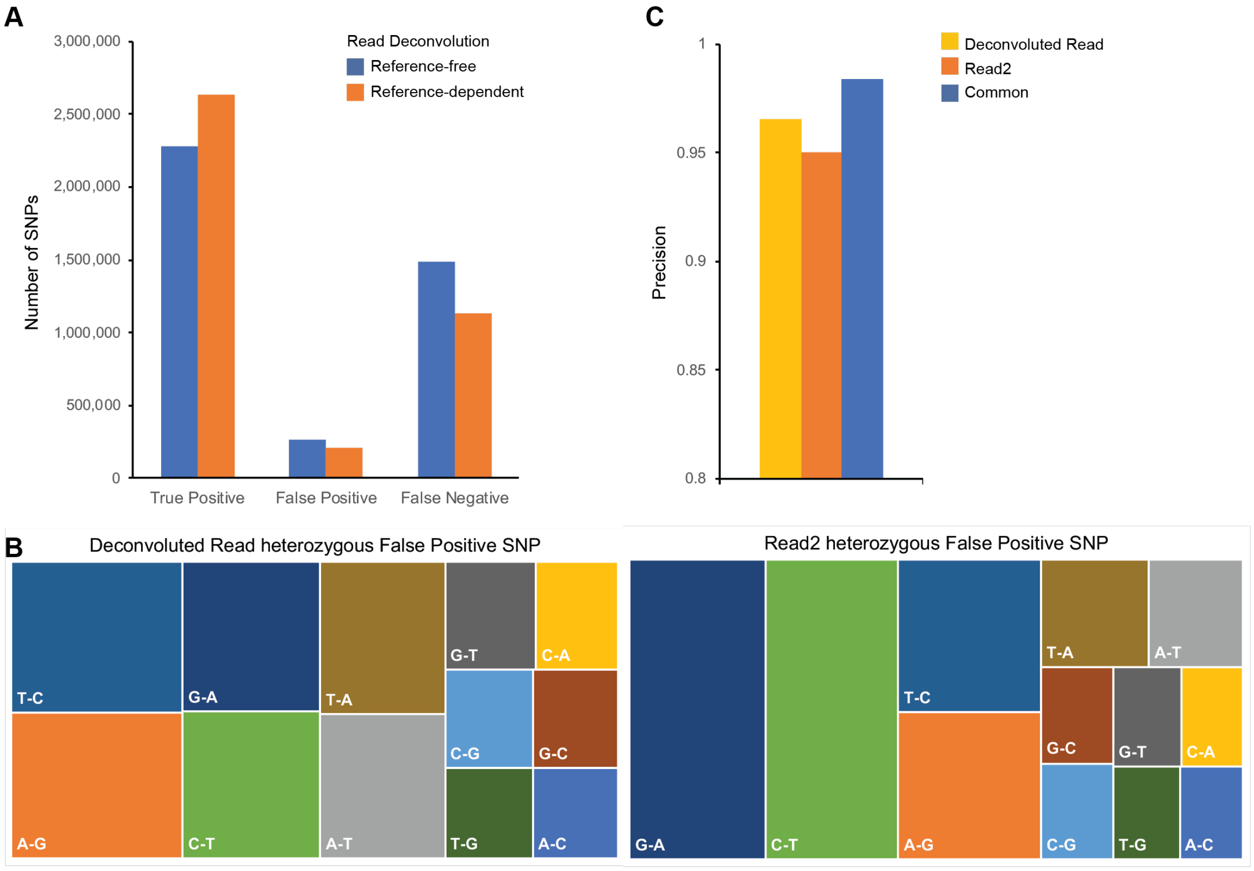


Supplemental Figure 2: Comparison of SNP calling with different strategies.
The defined SNPs were benchmarked against JIMB WGS data. (A) Methyl-SNP-seq Replicate 1 data was used for SNP calling using: Reference-free Deconvoluted Read and Reference-dependent Deconvoluted Read. (B) Characterization of the false positive heterozygous SNPs defined by Deconvoluted Read or Read2. (i.e. T-C means in the vcf file REF=T while ALT=C). (C) Precision of SNPs defined using Deconvoluted Read and Read2. The common SNPs are those detected by both Deconvoluted Read and Read2.

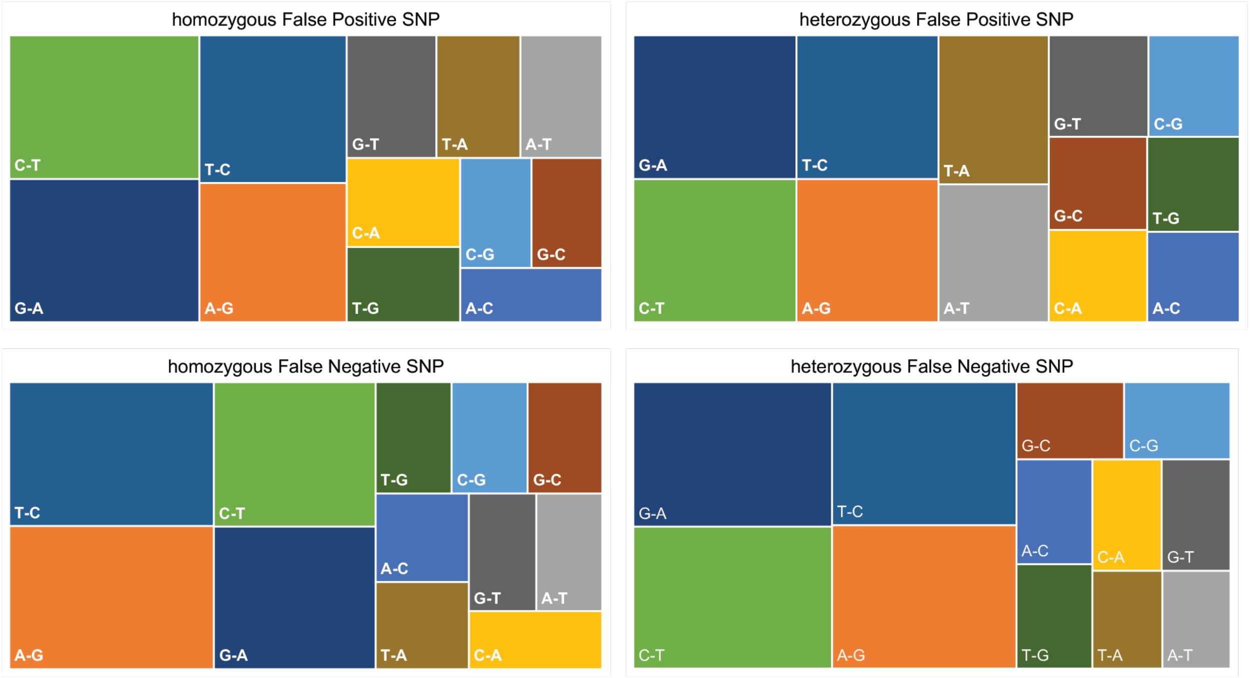


Supplemental Figure 3: Characterization of the Methyl-SNP-seq defined False positive or negative SNPs. The Methyl-SNP-seq defined SNPs were benchmarked against JIMB WGS defined SNPs. (i.e. T-C means in the vcf file REF=T while ALT=C).

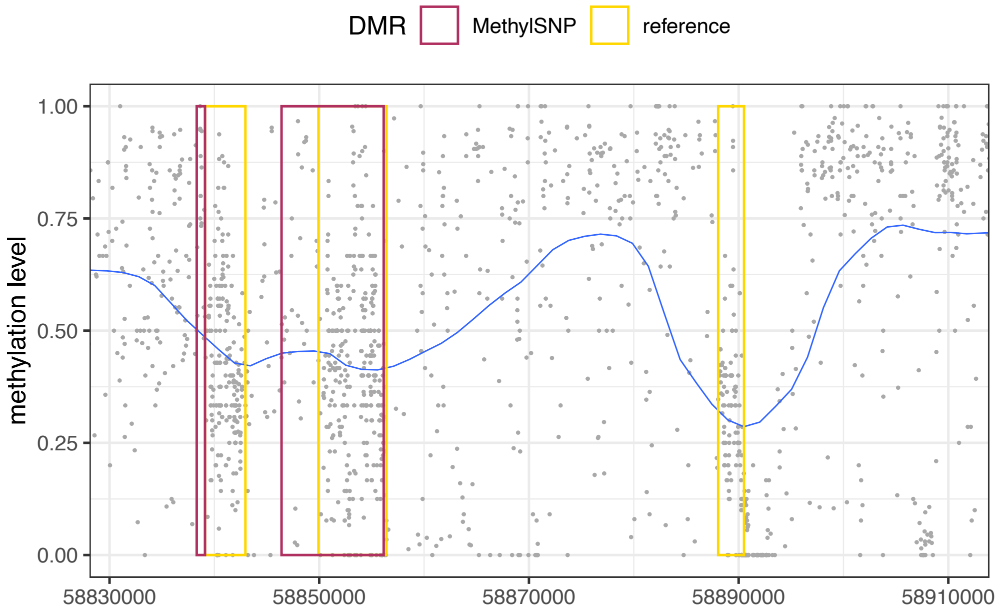


Supplemental Figure 4: CpG methylation profile and DMRs near the imprinted GNAS gene. Coordinates on the x axis correspond to positions on chr20 (GRCh38). Red box: Methyl-SNP-seq detected DMRs (the right 2 DMRs in Figure 4B examples). Yellow box: reported DMRs.

**Supplemental Tables**

Supplemental Table 1: Picard AlignmentSummaryMetrics

Supplemental Table 2: Picard VariantCallingMetrics

Supplemental Table 3: Picard GCBiasMetrics

Supplemental Table 4: Methyl-SNP-seq reported allele-specific DMRs overlap with known imprinted clusters and associated ASDMRs.
Methyl-SNP-seq reported allele-specific DMRs which overlap or near (within the window of upstream 10kb to downstream 10bk) the known allele specific DMRs (Fang et al. 2012) are listed in this table.

Supplemental Table 5: Association of ASDMRs with different genomic features

| Genomic regions | Biological Significance | Assay type and data source/reference | overlapping with ASDMR | overlapping with random CpG regions | Fold change |
|---|---|---|---|---|---|
| open chromatin | open chromatin | ATA-seq (ENCSR637XSC) | 3448 | 3650 | 0.9 |
| H3K27Ac | active enhancer | ChIP-seq (ENCSR000AKC (Creyghton et al. 2010)) | 1529 | 1773 | 0.9 |
| H3K4me1 | enriched at active and primed enhancers | ChIP-seq (ENCSR000AKF (Rada-Iglesias 2018)) | 2397 | 1887 | 1.3 |
| H3K4me1 - H3K4me3 | general enhancers including active and primed enhancers, non-promoters | (Creyghton et al. 2010) | 1594 | 1090 | 1.5 |
| H3K4me3 | promoters and active transcription | ChIP-seq (ENCSR057BWO (Mikkelsen et al. 2007)) | 1142 | 1716 | 0.7 |
| H3K9me3 | heterochromatin; frequently coexist with methylation | ChIP-seq (ENCSR000AOX) | 513 | 355 | 1.4 |

Supplemental Table 6: Picard target enrichment HsMetrics for targeted Methyl-SNP-seq

### Supplemental Protocol

Buffer used:

LowTE: 1mM Tris pH8.0 and 0.1mM EDTA

Preparation of 20uM annealed hairpin adapter:
10X annealing buffer: 100mM Tris pH8.0 and 1M NaCl
Dilute the hairpin adapter to 20uM in 1X annealing buffer. Anneal as follows: incubate at 80 degree for 2 min then ramp temperature to 25 degree at a rate of 0.1 degree per second. The annealed hairpin adapter is saved at -20 degree.

Preparation of 50ul 10mM d(A,T,G)TP mix:
100mM dATP 5ul + 100mM dGTP 5ul + 100mM dTTP 5ul + 35ul H2O

1. Fragmentation of genomic DNA (gDNA)

Sonicate 2ug NA12878 gDNA (add diluted non-methylated lambda gDNA as spike-in) in 50ul 10mM Tris pH8.0 using Covaris S2 with the following setup:
250bp protocol, Duty Cycle: 10%, Intensity: 5, Cycles/burst 200, Time 80s.
Set up two sonication (total 4ug gDNA) for the human whole genome Methyl-SNP-seq sequencing.

2. End Repair and dA tailing using Ultra II End Repair/dA tailing module (NEB E7546)

| | |
|---|---|
| 4ug Fragmented gDNA | 100ul |
| NEBNext Ultra II End Prep Enzyme Mix | 9ul |
| NEBNext Ultra II End Prep Reaction Buffer | 21ul |
| H2O | 50ul |
| Total | 180ul |

- Incubate at 20 degree for 30 min, then at 65 degree for 30 min, then cool down to 12 degree.
- Clean up the 180ul reaction using zymo oligo clean (zymo D4060) with the 80nt protocol. Elute the column with 26ul LowTE.

3. Ligation to hairpin adapter and removal of unligated DNA

| | |
|---|---|
| dA tailed DNA from step 2 | 26ul |
| 20uM annealed hairpin adapter | 4ul |
| 2X Ligation Mix (NEB M0367) | 30ul |
| Total | 60ul |

- Incubate at 20 degree for 1 h.
- Clean up the 60ul reaction using 0.9X=55ul SPRI beads, elute with 40ul LowTE.

Then:

| | |
|---|---|
| Ligated DNA | 40ul |
| ExoIII (100U/ul) (NEB M0206) | 2.5ul |
| ExoVII (10U/ul) (NEB M0379) | 2.5ul |
| 10X NEB Cutsmart buffer | 5ul |
| Total | 50ul |

- Incubate at 37 degree for 1 h.
- Clean up the 50ul reaction using zymo oligo clean (zymo D4060) with the 80nt protocol. Elute the column with 40ul LowTE. Stop point: save product at -20 degree.

4. Nick translation and dA tailing

| | |
|---|---|
| Ligated DNA from step 3 | 39ul |
| 10X NEB Buffer 2 | 5ul |
| UDG (5U/ul) (NEB M0280) | 2ul |
| Endo IV (10U/ul) (M0304) | 2ul |
| Total | 48ul |

- Incubate at 37 degree for 15 min.

Then add directly to the reaction without purification:

| | |
|---|---|
| 10mM d(A,T,G)TP mix | 0.5ul |
| 10mM 5-methyl-dCTP (NEB N0356) | 0.5ul |
| NEB DNA pol I (10U/ul) (NEB M0209) | 1ul |
| Total | 50ul |

- Incubate at 20 degree for 30min.
- To add dA tail, add directly to the reaction without purification: 1ul 100mM dATP (final concertation 2mM), 2ul Taq polymerase (NEB M0267), incubate at 65 degree for 30min then cool down to 12 degree.
- Clean up the 50ul reaction using zymo oligo clean (zymo D4060) with the 80nt protocol. Elute the column with 14ul 10mM Tris pH8.0.

5. Ligation to methylated illlumina adapter

| | |
|---|---|
| dA tailed mC incorporated DNA from step 4 | 14ul |
| 7.5uM NEB EM-seq adapter | 1ul |
| 2X Ligation Mix (NEB M0367) | 15ul |
| Total | 30ul |

Note: Dilute the 15uM NEB EM-seq adapter (NEB E7165 included in the E7140 kit) using 10mM Tris pH8.0 or NEB adapter dilution buffer.

- Incubate at 20 degree for 1 h.
- Add volume to 50ul and clean up using 0.8X=40ul SPRI select beads. Elute with 20ul LowTE.

6. Bisulfite treatment

Perform the bisulfite conversion using Fast Bisulfite Conversion kit (Abcam ab11727) following the manufacture's instruction with the following modifications:
Evaporate the 20ul DNA from step 5 into 5ul to mix with the bisulfite reagent.
Incubate the bisulfite conversion reaction at 95 degree for 25 min.

7. PCR

Perform PCR amplification using NEBNext Q5U Master Mix (NEB M0597) following the manufacture's instruction.
In our study, we used 8 PCR cycles and got about 50ul 3nM-4nM amplified library, which was enough for at least 2 Novaseq runs.
